# Developmental Changes in Movement Related Brain Activity in Early Childhood

**DOI:** 10.1101/531905

**Authors:** Blake Johnson, Cecilia Jobst, Rita Al-Loos, Wei He, Douglas Cheyne

## Abstract

In a previous MEG study of movement-related brain activity in preschool age children, we reported that pre-movement fields and sensorimotor cortex oscillations differed from those typically observed in adults, suggesting that maturation of cortical motor networks is still incomplete by late preschool age (Cheyne et al., 2014). Here we describe the same measurements in an older group of school-aged children (6 to 8 years old) and an adult control group, in addition to repeated recordings in seven children from the original study approximately two years later. Differences were observed both longitudinally within children and between age groups. Pre-movement (readiness) fields were still not present in the oldest children, however both frequency and magnitude of movement-related mu (8-12Hz) and beta (15-30Hz) oscillations demonstrated linear increases with age. In contrast, movement-evoked gamma synchronization demonstrated a step-like transition from low (30-50 Hz) to high (70-90 Hz) narrow-band oscillations, and this occurred at different ages in different children. These data provide novel evidence of linear and non-linear changes in motor cortex oscillations and delayed development of the readiness field throughout early childhood. Individual children showed large differences in maturation of movement-related brain activity, possibly reflecting differing rates of motor development.

## 1. Introduction

The acquisition of skilled motor abilities is one of the most important aspects of human development, with many motor and cognitive abilities emerging during early childhood. Difficulties in acquiring motor skills often first appear during preschool age and may persist throughout adulthood if not resolved within the first decade of life (Smith 2006, Piek et al. 2012) and may be co-morbid with pervasive developmental disorders of language and cognition (McCleery et al. 2013). Surprisingly, little is known about the relationship between brain development and the acquisition of motor abilities in early childhood, even though human neuroimaging studies have demonstrated rapid changes in brain structure and function during this period (Durston and Casey 2006, Shaw et al. 2008, Brown and Jernigan 2012). We also know from observation that a child’s movements are not the same as an adult. Fine motor control and dexterity do not resemble those of the adult until late childhood and the development of these skills proceeds at different rates in different children (Forssberg et al. 1991, Dayanidhi et al. 2013). This is accompanied by notable differences in movement-related brain potentials. For example, Electroencephalography (EEG) measures of movement-locked brain potentials do not show a typical slow pre-movement negative shift – or “readiness potential” – until children reach 8 to 9 years of age (Chisholm and Karrer 1988, Chiarenza et al. 1995). Cortical oscillatory activity also demonstrates amplitude and frequency changes throughout childhood, likely reflecting changes in underlying sensorimotor networks (Southgate et al. 2009, Berchicci et al. 2011, Marshall et al. 2011, Thorpe et al. 2016, Trevarrow et al. 2018).

Magnetoencephalographic (MEG) recordings provide a well-established technique for studying the time course of brain activity during motor tasks, as well as the role of oscillatory networks in motor control (for review see, Cheyne 2013). With the introduction of customized MEG systems for smaller children, MEG can fill the current gap between our knowledge of early neural development and that of motor and cognitive abilities (Johnson et al. 2010, Cheyne et al. 2014, Roberts et al. 2014, He et al. 2015, Okada et al. 2016, He and Johnson 2018, Johnson and He in press). There are however, relatively few MEG studies of motor development, partly due to the challenges of recording motor responses in children and most studies to date have focused on older children and adolescents (Gaetz et al. 2010, Wilson et al. 2010, Huo et al. 2011, Trevarrow et al. 2018). In a previous study (Cheyne et al. 2014) we reported the first recordings of movement-related brain activity in preschool age children using a customized pediatric MEG system (Johnson et al. 2010). Using a child-friendly motor task and advanced source analysis techniques we were able to successfully localize movement-related brain activity to the sensorimotor cortex in all children, however as predicted, preschool aged children showed marked differences from the adult pattern of pre-movement waveform morphology, with an absent early readiness field component. Differences in both the timing and frequency and strength of motor cortex oscillations to those typically observed in adults were also observed, although some of the youngest children (less than 4 years of age) demonstrated completely adult-like patterns of high frequency motor gamma oscillations, suggesting that there are large individual differences in movement-related brain activity in preschool age children.

Without longitudinal measurements within the same children, it is unclear whether the individual differences observed in our earlier study may have persisted throughout development, or simply reflected differences in the rate of emergence of these brain responses across individuals. This has important implications for studies of normal motor development, as well the use of these measures as indicators of atypical development. Here we report a follow-up to our previous study of MEG recordings of motor activity in children, comparing data from preschool age children to recordings in an older group of children two to two and one-half years older to test the hypothesis that children showing less mature movement-related brain activity at preschool age would show a transition to the adult form. In addition, approximately half of the children in the older age group also participated in our first study, allowing us to observe such changes longitudinally within individual children. We also included a group of adults who performed the same motor task to confirm that differences are not task related. To our knowledge, this is the first longitudinal MEG data examining changes in neural activity in the motor cortex throughout early childhood.

## 2. Materials and Methods

### 2.1 Participants

We conducted MEG recordings of movement-related brain activity in children from two age groups in two different recording sessions (details provided in Table 1). The first group (“**Session 1**”) consisted of fifteen healthy preschool age children (8 female, mean age 4.20 years ± 0.19, range 3.2 to 4.9 years old, all right-handed) who were included in our previously published study (Cheyne et al. 2014). For the follow-up study (“**Session 2**”), we recruited a second group of school-aged children; seven from the original study who had been scanned approximately two years earlier and ten additional children matched for age and sex to the children from the first study. Four children were excluded from final analysis due to poor task compliance or excessively noisy data, leaving a total of 13 children (6 female) with a mean age of 6.79 ± 0.21 years in Session 2, ranging from 5.5 to 7.5 years old (referred to hereafter as 6 to 8-year-olds) with a subgroup of seven children in whom longitudinal changes could be directly examined. Thirteen adults (6 female, 1 left-handed, mean age 31.1 years, range 20.3 to 39.3 years) also participated in Session 2 to provide an adult comparison group performing the same motor task. Handedness in adults was assessed using the Edinburgh handedness inventory (Oldfield 1971). Handedness was assessed in the younger children by observing which hand they used to perform a number of simple actions (hammering, writing their name or drawing a picture, opening a jar, putting crayons into a box, throwing a ball and cutting a piece of paper with child-safe scissors). Participants were recruited from the Sydney area and provided informed consent using protocols approved by the Macquarie University Human Subjects Ethics Committee.

**Table 1.** Fractionized age, sex and handedness of participants in each age group, adult, preschool and school-age. Bold font indicates children imaged in both sessions 1 and 2.

| Session 1: Preschool |  |  |  | Session 2: School Age |  |  |  | Healthy Adults |  |  |  |
| --- | --- | --- | --- | --- | --- | --- | --- | --- | --- | --- | --- |
| *Subject* | *Sex* | *Handedness* | *Fractional Age* | *Subject* | *Sex* | *Handedness* | *Fractional Age* | *Subject* | *Sex* | *Handedness* | *Fractional Age* |
| **0493** | M | Right | **3.93** | **0493** | M | Right | **5.81** | 1662 | M | Right | 29.82 |
| **0494** | M | Right | **3.68** | **0494** | M | Right | **6.34** | 1669 | M | Right | 31.38 |
| **0498** | M | Right | **4.01** | **0498** | M | Right | **6.50** | 1670 | M | Right | 33.65 |
| **0499** | M | Right | **4.61** | **0499** | M | Right | **7.27** | 1676 | F | Right | 31.85 |
| **0511** | F | Right | **4.84** | **0511** | F | Right | **7.36** | 1677 | F | Right | 27.16 |
| **0514** | F | Right | **4.72** | **0514** | F | Right | **7.25** | 1680 | F | Right | 35.00 |
| **0515** | M | Right | **3.64** | **0515** | M | Right | **6.17** | 1685 | M | Right | 28.53 |
| 0501 | M | Right | 4.60 | 1674 | M | Right | 7.73 | 1688 | F | **Left** | 26.42 |
| 0502 | F | Right | 4.70 | 1675 | F | Right | 7.13 | 1703 | F | Right | 24.18 |
| 0503 | F | Right | 4.02 | 1692 | F | Right | 6.98 | 1707 | F | Right | 20.27 |
| 0505 | M | Right | 4.30 | 1693 | F | Right | 7.48 | 1711 | M | Right | 32.71 |
| 0506 | F | Right | 3.16 | 1718 | F | Right | 5.46 | 1721 | M | Right | 25.45 |
| 0509 | F | Right | 4.39 |  |  |  |  | 1727 | M | Right | 39.28 |
| 0510 | F | Right | 4.41 |  |  |  |  |  |  |  |  |
| 0536 | F | Right | 4.86 |  |  |  |  |  |  |  |  |
| *Mean Age*<br>*+/- SEM*<br>*(Range)* |  |  | *4.20 +/- 0.19*<br>*(3.64 - 4.84)* |  |  |  | *6.79 +/- 0.21*<br>*(5.46 - 7.73)* |  |  |  | *31.06 +/- 1.05*<br>*(27.16 - 35.00)* |

### 2.2 Motor Task

The same videogame-like task was used in both recording sessions to elicit self-paced motor responses. Children were instructed to press a button when specific images appeared on a display in a space-theme game. Images of space aliens appeared at the top of a screen every 3 s and descended slowly towards a space ship at the bottom of the screen. Participants were instructed to “feed the aliens” as they appeared by pressing a button with their right or left index finger, which would launch cookies from the space ship towards the approaching aliens. Participants were also coached to maintain their own timing to encourage more self-paced movements as in our previous study. Children performed the task for approximately 5 minutes each for left or right-hand responses (order counter-balanced across participants), which was sufficient in obtaining approximately 100 button responses for each hand. To help keep the children engaged in the task, explosive sounds were heard when the cookie reached the target; however, these were not time-locked to the appearance of the stimulus or the button press. Adults performed the task with their right hand only. Motor responses were measured with a response button pad (LUMItouch Response System, Lightwave Medical Industries, Burnaby, Canada). Visual stimuli were presented on a back-projected screen mounted approximately 140 cm above the subject using an InFocus LCD projector (Model IN5108). Auditory feedback was presented via plastic tubes with insert earphones (Etymotic Research Inc., Model E-30, Elk Grove Village, IL).

### 2.3 Data Acquisition

MEG data in the children was acquired with a custom, child-sized MEG system (Model PQ1064R-N2m, Kanazawa Institute of Technology (KIT), Kanazawa, Japan) in a magnetically shielded room (MSR) (Fujihara Co. Ltd., Tokyo, Japan) located at Macquarie University. The system consisted of 64 first-order axial gradiometers and is designed to fit approximately 90% of heads of 5-year old children (Johnson et al. 2010). Session 2 scans were collected on the same child system, although approximately half of the participants were measured after the sensor array was upgraded from 64 to 125 sensing channels. Adults were measured in a 160-channel adult MEG (Model PQ1160R-N2, KIT, Kanazawa, Japan) located within the same magnetically shielded room. All data was acquired at a sampling rate of 1000 samples per second with a 0.3 Hz to 200 Hz bandwidth. All participants were recorded in the supine position with button pad positioned on their chest.

Head shape data was collected with a pen digitizer (Polhemus Fastback, Colchester, VT) for all participants, which included the digitization of three fiducial points on the head (nasion and pre-auricular points), as well as five head position indicator (HPI) coils that were secured to the head by an elasticized head cap that tracked total movement (pre and post MEG recording head location measurements are averaged together). Off-line sensor data was transformed into a head-based coordinate system based on the digitized fiducial points and HPI coils.

### 2.4 Data Analysis

All MEG data analyses and visualization was carried out using the *BrainWave* toolbox (Jobst et al. 2018) developed at the Hospital for Sick Children implemented in C++ and *MATLAB* (Mathworks, Natick, MA). The continuous KIT (.con) data recordings were imported into *BrainWave* and segmented into 4 second trials (2 second prior to and 2 seconds following the first button press in response to the target appearance). Each trial was examined for large signal deviations (eye-blinks, movement artefacts) and removed prior to further analysis. In order to avoid data processing differences between Session 1 and Session 2 all data from the first session was reanalyzed using the identical analysis steps.

As in our previous study, a paediatric MRI template customized for children 4.5 to 8.5 years of age from the NIHPD database (Fonov et al. 2011) was warped to fit each participant’s digitized head shape using the iterative closest point (ICP) algorithm, producing a unique “surrogate” MRI for each participant for spatial normalization of source images to a standard (MNI) template brain using *SPM8* (http://www.fil.ion.ucl.ac.uk/spm/software/spm8/). As shown in our previous study, this approach provides a good approximation to MNI coordinate space for group averaging of whole-brain source images and for obtaining anatomical labels (Brodmann areas). Surrogate MRIs were similarly created for adults using the Colin-27 adult template MRI (Holmes et al. 1998). MEG source modeling was carried out using a single-sphere head model (Sarvas 1987) fit to the inner skull surface extracted from the (warped) surrogate MRI (Lalancette et al. 2011). The inner skull surface was extracted using *FSL* (http://fsl.fmrib.ox.ac.uk/fsl/fslwiki/).

Movement evoked activity was localized for a 600 ms window around time zero (−0.3 s to 0.3 s in time increments of 5 ms) averaged across trials using a wide-band 1-30 Hz event-related beamformer (ERB) algorithm (Cheyne et al. 2006, Cheyne et al. 2007) implemented in the *BrainWave* toolbox. Beamformer weights were calculated using the data covariance −0.5 s to 1 s around the time of the button press (t = 0.0), with diagonal regularization of 10 femtoTesla/sqrt (Hz). For visualization of source activity, individual and group averaged source images were interpolated onto an inflated cortical surface extracted from the template MRI used for spatial normalization using the *FreeSurfer* toolbox (http://surfer.nmr.mgh.harvard.edu/).

### 2.5 Statistical Analyses

Time courses of neural activity were compared for the different age groups using the method described by Guthrie and Buchwald to estimate the minimum number of empirically significant consecutive paired t-tests between the two time series. Implemented in a MATLAB toolbox (Siegle et al. 2003), this method uses the pooled autocorrelation value to threshold the minimum number of consecutively significant t-values representing a p-value of less than 0.05 within 150 ms time windows clusters along the entire time series (Guthrie and Buchwald 1991).

Movement-induced power change in specific frequency band ranges of interest; mu (7-13 Hz), beta (15-30 Hz), low gamma (30-60 Hz) and high gamma (60-90 Hz) using a Morlet wavelet-based time-frequency decomposition implemented in the *BrainWave* toolbox. Time-frequency representation (TFR) plots were computed using 5 cycles per wavelet for the mu frequency band (9 cycles for beta and 20 cycles for low and high gamma) and transformed to percent change from a baseline period of −2 s to 0 s preceding the button press. Averaged (phase-locked) source power was subtracted from the single-trial power for each individual to remove any power changes related to the evoked response. Peak frequency and amplitude within a specified frequency band and time window were estimated from the TFRs using a previously described bootstrapping procedure (Muthukumaraswamy et al. 2009, Cheyne and Ferrari 2013).

## 3. Results

### 3.1 Movement-related Fields

Averaged time courses of peak activations in the event-related MEG source images revealed typical motor responses in individual subjects localized to the contralateral hand region of sensorimotor cortex (BA 4/6) around the time of movement onset (button press). Since the children did not have a pre-movement readiness or motor field peak, the first peak of activity immediately following movement onset was used as to extract their sensorimotor activation time course. Figure 1 shows the group averaged source activity peaking approximately 20 ms after movement onset for right hand button press for adults and both child age groups (Session 1 and Session 2), projected onto the inflated surface of the MNI template brain showing activation in the left sensorimotor hand area and corresponding group averaged source activity waveforms. Time courses of source activity for each individual were computed using the location of their individual peak activation. Negative deflections in all plots reflect anterior directed dipole sources in MI after correcting for polarity reversals across subjects.

**Figure 1.**
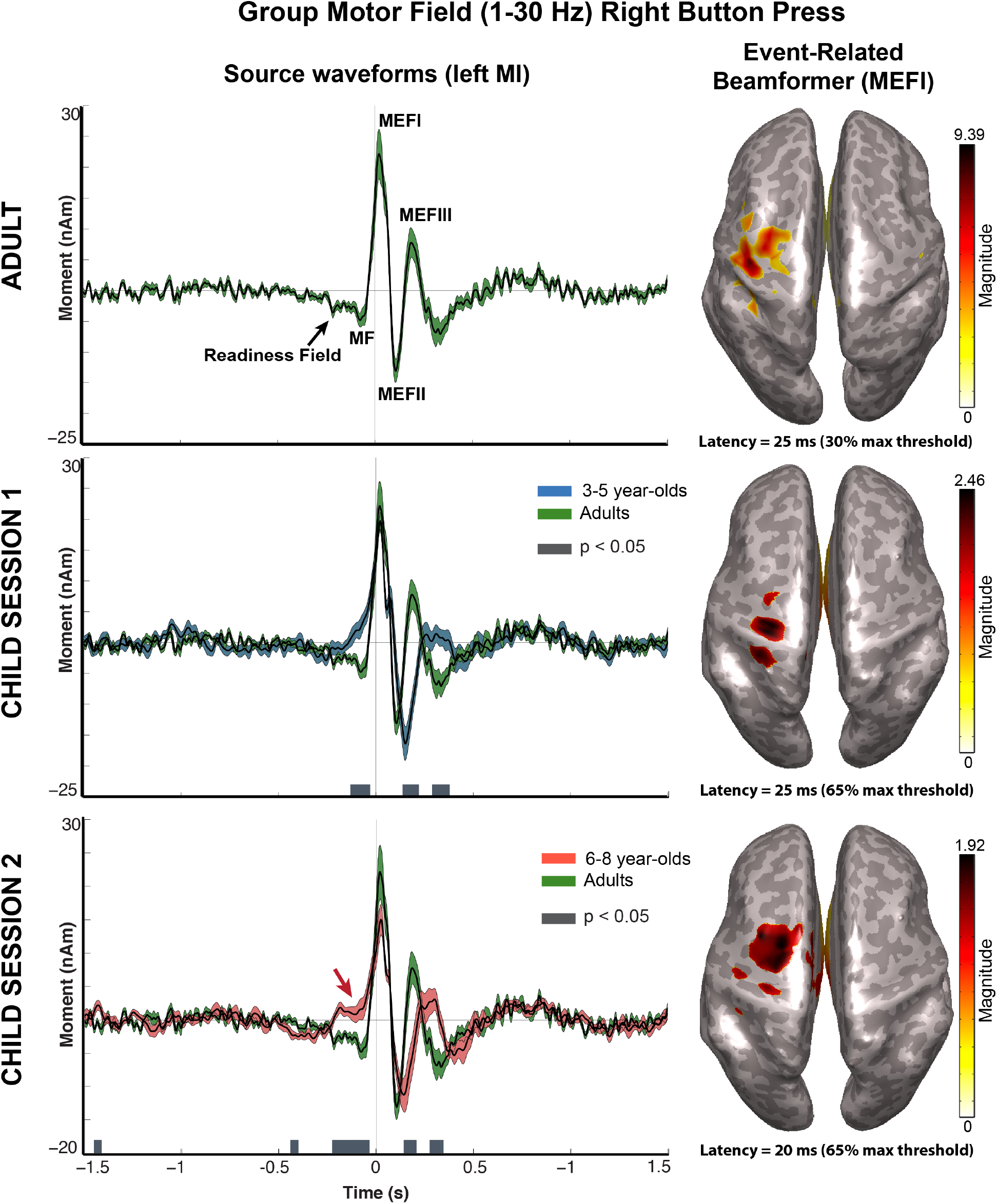
Group averaged source activity in the contralateral sensorimotor cortex time-locked to right index finger movements (button press) for adults and children. The colour scale reflects source strength (in units of nanoAmpere-meters) at the latency of the MEFI at 20 to 25 ms following movement onset are projected onto an inflated template brain (the Colin-27 adult template for adults, and a child template for 4.5-8.5 year-olds for children), thresholded at 30-65% of the group maximum. Source waveforms computed for peak activity closest to the primary motor cortex in individual subjects is shown on the left. Adult source activity (top row) shows typical premovement readiness field, peaking with a motor field (MF) at approximately 50 ms prior to button press followed by movement-evoked field components (MEFI-III). Source waveforms for Session 1 (3-5 year-olds) and Session 2 (6-8 year-olds) are shown bel; superimposed on the adult waveforms. Shading represents the standard error across subjects. Grey horizontal bars indicate periods during which waveforms are significantly different (p<0.05). The emergence of a motor field peak is apparent in the older children (red arrow) along with delayed MEF components.

Source waveforms in the adults demonstrated a typical slow shift or readiness field beginning approximately 500 ms prior to movement, peaking approximately 50 ms prior to the button press (motor field, MF), followed by three field reversals or “movement-evoked” fields (MEFI, MEFII and MEFIII) peaking at approximately 20 ms, 100 ms and 190 ms after the button press, respectively. Periods of significant amplitude differences between waveforms (using Guthrie-Buchwald correction at p < 0.05) are indicated by horizontal grey bars. There was a clear absence of a pre-movement MF peak prior to movement onset in both child age groups as well as amplitude differences during the movement evoked fields, due to an apparent earlier peak latency of the MEFII in the adults. The latter was confirmed by comparing MEFII latency in adults which was significantly earlier (112 ms +/− 0.004 s.e.m.) compared to preschool (160 ms +/− 0.010 s.e.m., p = 0.0001) and school age children (135 ms +/− 0.012 s.e.m., p = 0.0096).

Amplitude differences between the young and old child groups were not statistically significant prior to or following movement onset. As shown in Figure 2, movement-related waveforms were remarkably similar for left and right movements in both age groups. In comparison to the younger age group, a small deflection around 200 ms prior to movement onset was apparent in the 6 to 8-year-olds (also shown in Fig. 1, lower row) for both left and right movements, suggesting the emergence of a pre-movement motor field, although this was not statistically significant in the group data. In addition, there was a shift of the MEFII peak to an earlier latency in the older children in the direction of an expected adult response, however this was not statistically significant between the two child groups (*p* = 0.075).

**Figure 2.**
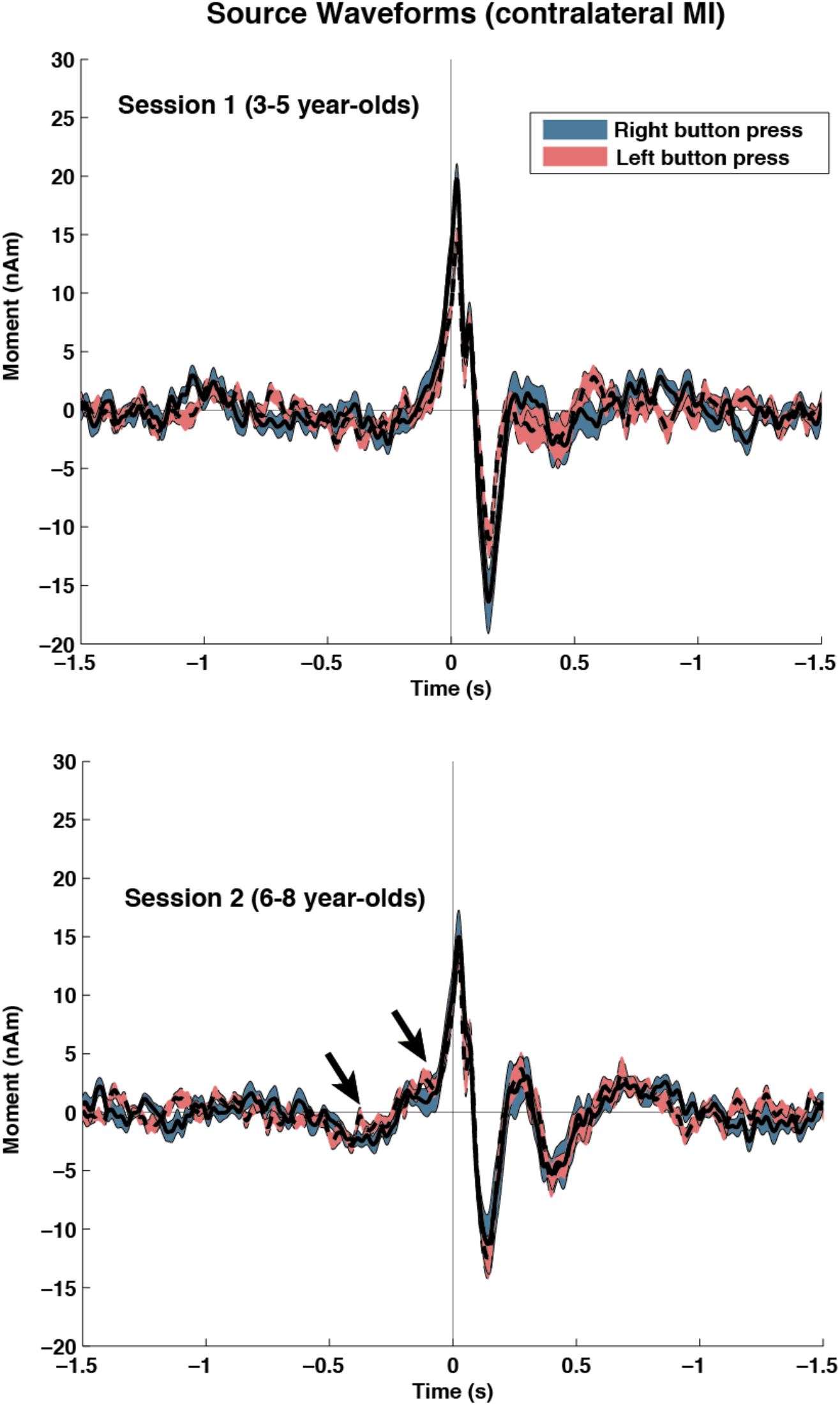
Group source waveforms for sensorimotor cortex sources superimposed for left and right button press conditions plotted separately for Session 1 (3-5 year-olds) (top) and Session 2 (6-8 year-olds) showing high similarity between left and right movements. A small deflection can be seen in the older child group at the latency of the motor field 50 to 100 ms prior to movement onset.

### 3.2 Sensorimotor Mu and Beta Oscillations

Figure 3 shows a time-frequency representation plots of the modulation of source power in the contralateral sensorimotor cortex from both child groups. As described in our original study, preschool age children demonstrated a pre-movement suppression and post-movement increase (rebound) of power in the mu (7-13 Hz) and beta (15-30 Hz) frequency bands that is much weaker and delayed compared to that observed in adults. In comparison, the 6 to 8-year-old group in the current study demonstrated a more robust beta suppression and an earlier and stronger beta rebound. Beta rebound power was positively correlated with age in children pooled across both age groups for both left sided (Pearson’s *r* = 0.603, p = 0.0008) and right-sided (*r* = 0.504, p = 0.007) movements. Changes in oscillatory power with age were not consistent for any other frequency bands or time periods in the children. Beta rebound power in the adults was clearly earlier and more robust in comparison to the children, but also displayed greater variability in power and peak frequency across individuals (data not shown). We did not find any statistically significant differences in any of the above measures between male and female subjects in either the adult or child age groups.

**Figure 3.**
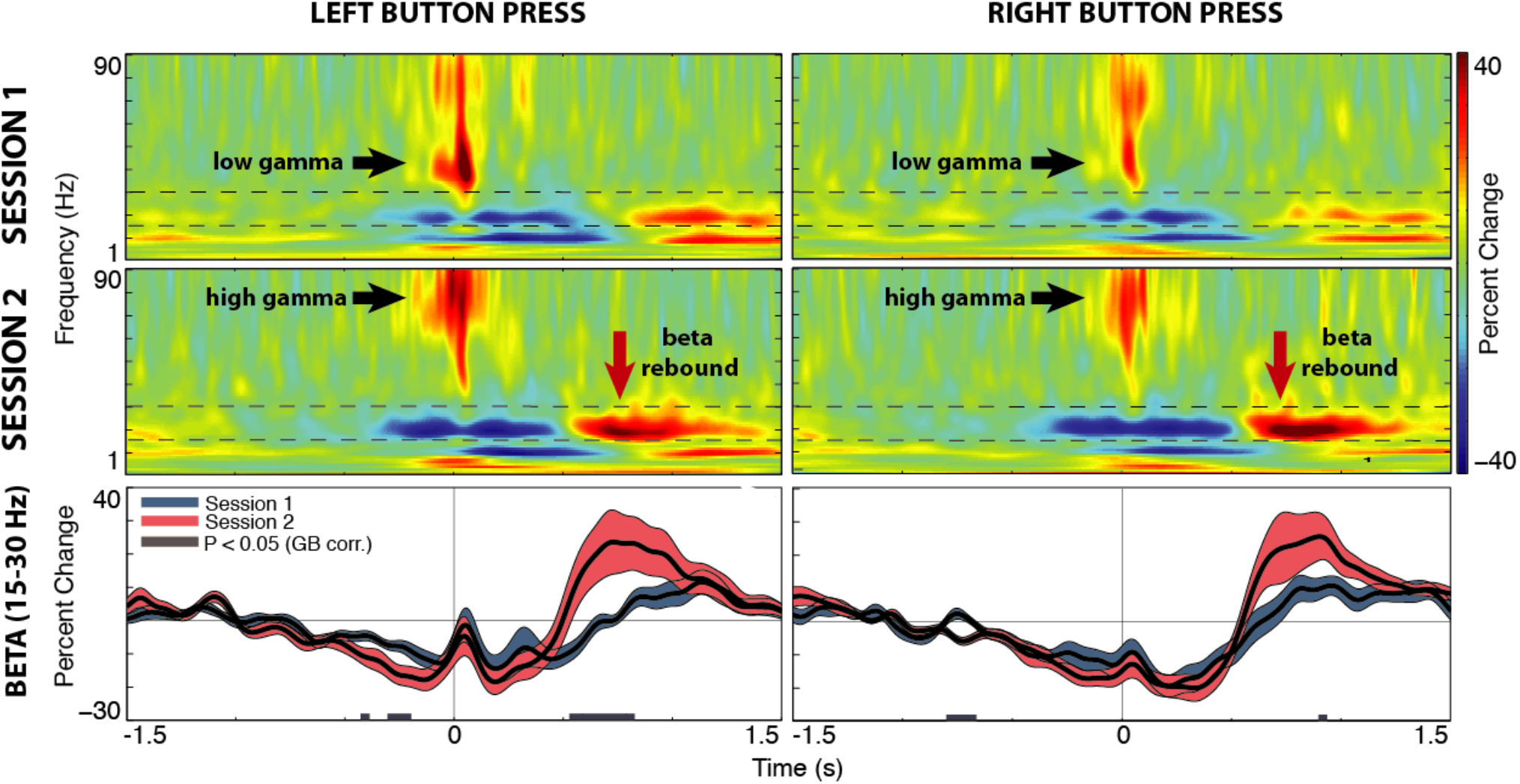
Time-frequency representation (TFR) plots of single-trial source power source power for Session 1 (3-5 year-olds) (top) and Session 2 (6-8 year-olds) age groups. Notice earlier and more robust beta activity and rebound and shift to high-frequency (70-90 Hz) gamma activity in the older children. Lower plot shows the time course of power averaged over the beta band (15-30 Hz) demarcated by the dotted lines in the TFR. Shading represents the standard error across subjects and periods of significant differences (*p* < 0.05) between the time series are indicated by horizontal gray bars.

To examine whether frequency of sensorimotor oscillations increased with age, mean peak frequency was estimated for the mu and beta bands for all age group during both pre-movement suppression (−200 ms to +700 ms) and post-movement rebound (+800 ms to +1600 ms) time windows, as shown in Figure 4a. Peak frequency of both mu and beta oscillations during premovement beta suppression significantly increased with age across all age groups (all comparisons using the Mann-Whitney test one-tailed t-test for unequal variances) for right button press but not for left button press (although mean frequency showed the same increasing trend). Mu peak frequency also increased during the rebound time window between children and adults, but was only significant between child age groups for left hand movements (p < 0.01). Peak frequency of beta rebound did not differ significantly across age groups with the exception of the young age groups versus adults, again for right hand movements only. Plotting peak mu and beta frequency pooled across all children and adults for right-hand movements as a function of fractionalized age (Figure 4b), revealed a significant linear increase of mu and beta frequency with age both during pre-movement suppression and post-movement rebound.

**Figure 4.**
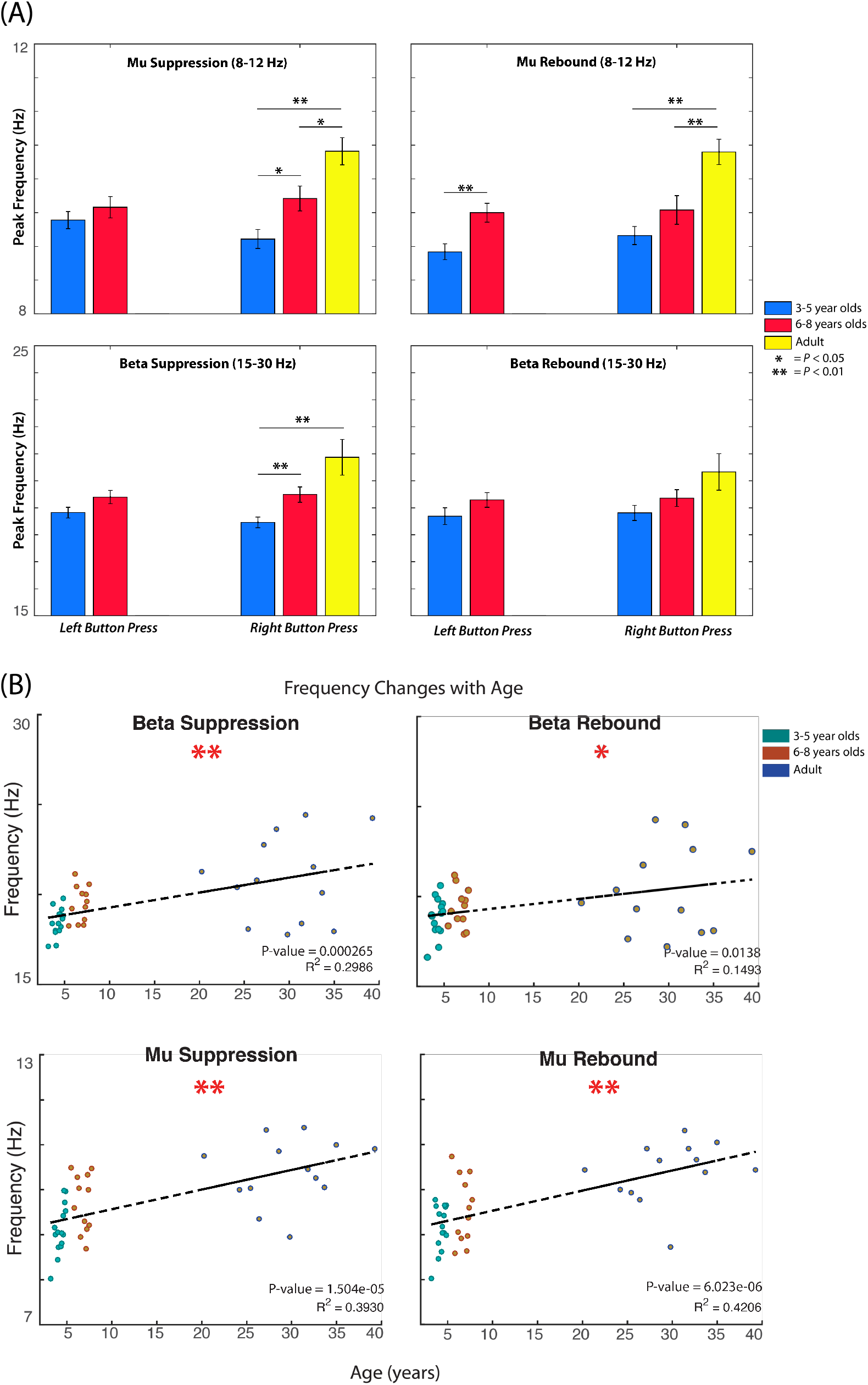
(A) Mean motor peak frequency (percent change relative to a baseline of −0.2s to 0s) is shown for mu and beta band oscillations during the left and right button press pre-movement suppression time window (−0.2 s to +0.7 s) and post-movement rebound time window (0.8 s to 1.6 s) for Session 1 (3-5 year-old), Session 2 (6-8 year-old) and compared with adults (right button press only). (B) Scatter plots of peak frequency of beta and mu band oscillations during the premovement suppression time window (left) and post-movement rebound time window (right) as a function of fractional age for all participants.

As reported in the first study, some preschool age children displayed a 70-90 Hz narrow-band gamma (also termed gamma event-related synchronization, ERS) at movement onset as previously reported in adults, while others had weak or absent movement-evoked gamma ERS or in some cases, a similar gamma ERS at movement onset, however at a much lower frequency (30-50 Hz). In contrast, all children in the 6 to 8-year-old cohort in Session 2 demonstrated a clear high frequency 70-90 Hz gamma burst (ranging from 50% to 150% above baseline) for both left and right movements, as can be seen in the group averaged TFR in Figure 3.

### 3.3 Longitudinal Changes between Session 1 and Session 2

Changes in both motor field patters and oscillatory activity were examined separately for the seven children who returned for Session 2 who were now 2 to 2 1/2 years older. Many of children recorded again in Session 2 showed highly similar, almost completely overlapping movement-related field patterns for both age time points. However, some children showed qualitative changes in their pre-movement source activity, as shown in Figure 5, with the emergence of a clearer MF and MEFI polarity reversal at movement onset. This likely contributes to the small deflection observed in the group source waveforms for both left and right movements shown in Figure. 2.

**Figure 5.**
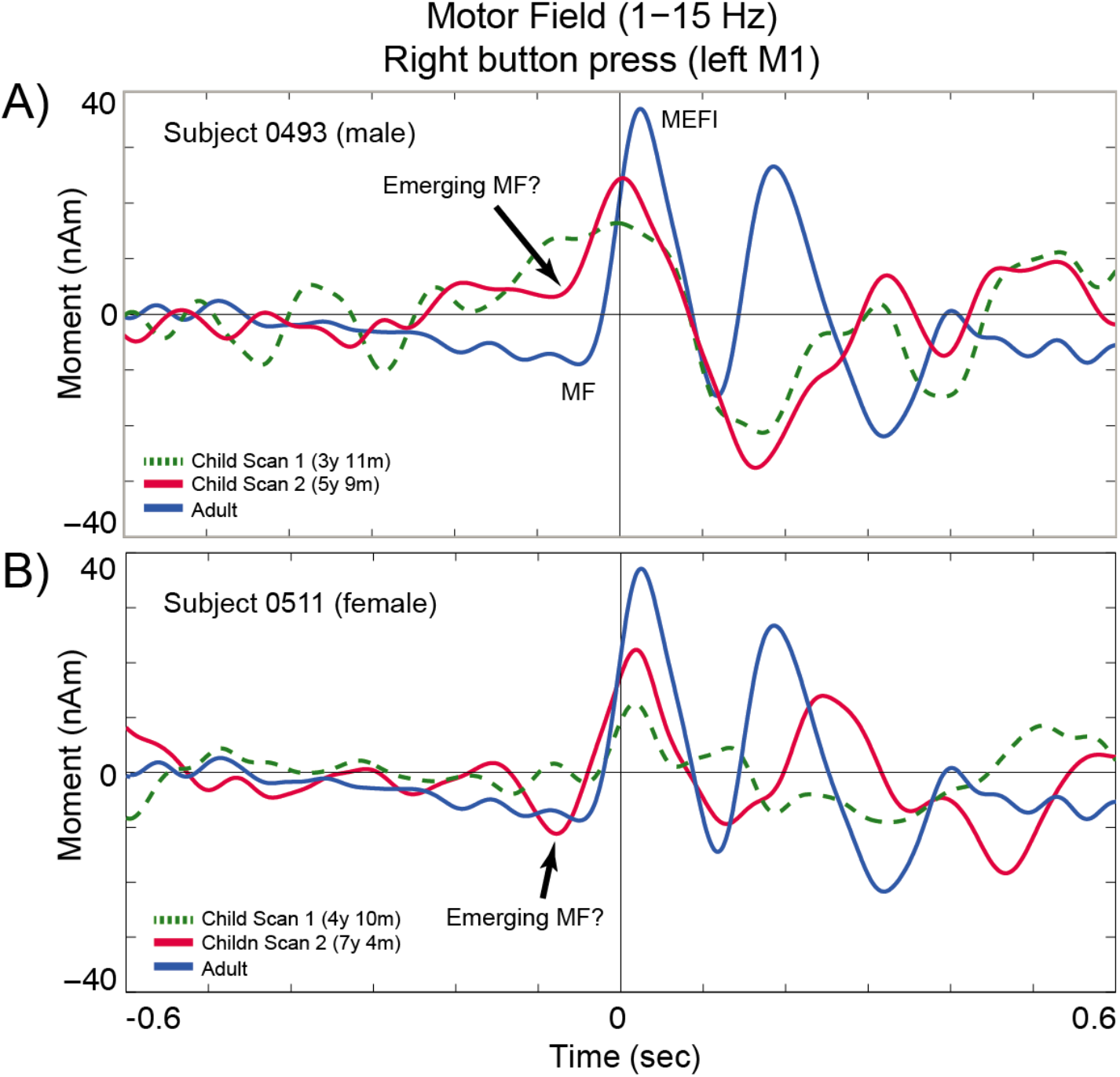
Longitudinal changes in movement-related source activity in two children measured at two time points, approximately 2 years apart (age at time of measurements is indicated in legend), compared to a typical response in one adult subject. Both children showed a weak motor field response in their first session (green dotted trace) but improved post-movement peaks (MEFI and MEFII) along with indications of an emerging pre-movement motor field (red trace) but no clear slow readiness field shift.

It was noted that children who did not show high frequency movement-evoked gamma in their first session, demonstrated a clear transition from low gamma burst at movement onset to high frequency gamma when comparing their Session 1 and Session 2 data. This was consistent for both left and right-hand movements as shown in Figure 6 (top row). In addition, children who did show a high frequency (adult-like) motor gamma pattern when measured at preschool age in Session 1 showed almost identical timing and mean frequency of gamma activity for both left and right movements when measured when two years older in Session 2 (Fig. 6, bottom row).

**Figure 6.**
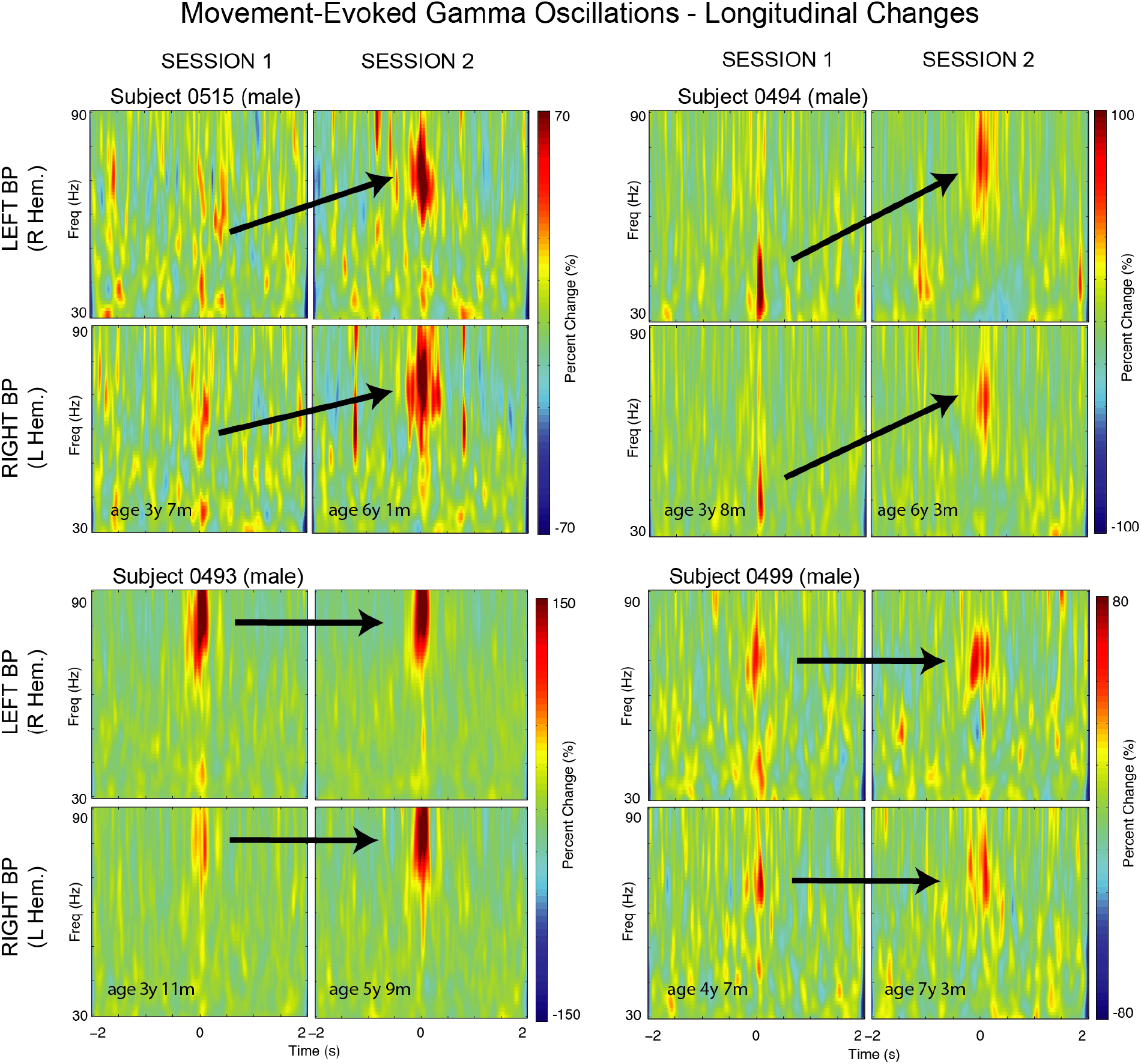
Examples of longitudinal changes of motor cortex gamma event-related synchronization (ERS) at movement onset in the gamma range (30-50 Hz). Top row: Two children showing transition (arrows) from a burst of low-frequency gamma ERS in the 30-50 Hz range (left) in their first session to the higher 70-90 Hz gamma burst in their second session (right) for both left and right movements. Bottom row: Two children who had clear high 70-90 Hz gamma ERS in both the first and second recordings sessions approximately two years apart. Note the high degree of similarity of gamma ERS strength and frequency for left and right movements within children.

## 4. Discussion

We examined movement-related fields and modulation of sensorimotor oscillations accompanying motor responses to visual cues in a video-game like task over two separate recording sessions in children ranging from 3 to 8 years of age compared to adults performing the same task. Subjects were instructed to respond to targets at their own pace to avoid strict time-locking to the visual stimulus. Results in the adult control group confirmed that our task elicited movement-related readiness fields. These were typical of those observed in healthy adults during self-paced movements, with pre-movement activity beginning about 1/2 second prior to movement reaching maximum amplitude just prior to a button press, corresponding approximately to EMG onset in the active muscles (Kristeva et al. 1991, Nagamine et al. 1996, Cheyne et al. 2006, Cheyne 2013). Movement-related brain activity in our participants was consistently localized to the hand area of the contralateral precentral gyrus (BA 4/6) confirming the validity of template MRI warping procedure for group averaging and obtaining approximate anatomical locations.

### 4.1 Age-related changes in movement-related fields

Motor responses shown for the preschool children demonstrate notable differences when compared to adults, as described in our previous study (Cheyne et al. 2014). However, even the oldest children in the 6 to 8-year-old cohort still did not demonstrate a mature slow readiness field or motor field peak reversal prior to the button press. Instead, pre-movement activity in children showed a rapid increase just prior to button press peaking at 20 ms following movement onset with the same polarity as the MEFI. This pattern was remarkably similar in some of children who returned up to 2.5 years following their initial recordings session, confirming that our task elicited stable and replicable motor responses, even in very young children. Movement-evoked fields also differed in children compared to adults, with longer MEFII and MEFIII peak latencies, which might be related to differences in movement duration or other kinematic parameters. It is interesting to note however that the MEFI component was present at a similar latency as adults. This movement-evoked brain response is thought to be related to proprioceptive feedback to the sensorimotor cortex during muscle contraction (Cheyne et al. 1997). In this regard, the comparable latencies of child and adult MEFI components is consistent with MEG responses to tactile stimulation of the hand in children, which show similar latencies to adults by 6 years of age (Pihko et al. 2009). While the afferent pathways in children are less myelinated and have slower conduction velocities, this is thought to be offset by the shorter travel distance in smaller arms (Pihko et al. 2009).

Our study included equal numbers of boys and girls, however there were no notable sex differences in the measured brain responses. This may reflect the fact that sex differences in the development of both gross and fine motor skills in typically developing children are relatively small (Flatters et al. 2014) and likely require a larger number of subjects to be detected.

The lack of a readiness field preceding movement in children, even in the oldest participants, remains puzzling given its ubiquitous presence in adults. Our results are consistent with EEG studies showing absent readiness potentials in children up to 9 years of age (Chisholm and Karrer 1988, Chiarenza et al. 1995). Although there were signs of the emergence of pre-movement field component, this was not statistically significant in the group data and likely reflects the large amount of variability observed within this age group. Additional studies in older children are required to determine when this important brain response develops and matures. However, our current findings support the notion that the readiness potential, thought to reflect motor preparation or intention (Cunnington et al. 2005, Colebatch 2007) is a late developing phenomenon, and raises questions regarding its role in motor control. Pre-movement activity is also observed in ipsilateral motor cortex in adults (Kristeva et al. 1990) and it has been proposed that this may reflect interhemispheric inhibition of contralateral motor cortex, particularly during the performance of skilled unilateral movements (Duque et al. 2007, Wischnewski et al. 2016). Notably, these inhibitory pathways do not appear to be fully developed until 10 years of age (Nezu et al. 1997) which roughly corresponds to the disappearance of mirror movements during development (Mayston et al. 1999). Although we were not able to assess the presence of mirror movements during the motor task in the current study, movement-related activity was largely contralateral to the side of movement, and differences were more significant for right hand compared to left hand movements in children who were assessed to be right-hand dominant, possibly reflecting a role of hand dominance in the maturation of movement-related brain activity.

### 4.2 Age-related changes in sensorimotor oscillations

In contrast to movement-related fields, we observed consistent differences between the two child age groups in sensorimotor oscillations, with a linear increase in mu and beta band peak frequency with age. We also observed significantly stronger and earlier onset of post-movement beta rebound in the 6 to 8-year-old group, indicating that beta rebound is robust by this age, although we cannot rule out that this might also reflect shorter duration movements in the older children, as beta rebound has been shown to be time-locked to movement offset rather than onset (Pfurtscheller et al. 1996, Cassim et al. 2001, Jurkiewicz et al. 2006) and our button box did not allow us to measure movement duration. We also observed a small but consistent increase in mu band frequency with age. In a review of mu rhythm activity across multiple age groups, Berchicci and colleagues (Berchicci et al. 2011) reported a rapid increase in mu frequency from infancy up to one year of age, followed by a more gradual increase in early childhood with lower (8-9 Hz) mean frequency compared to the 10 to 12 Hz range observed in adults, although no data were available for children older than 5 years of age. Our data complement these findings, showing a continued increase in mu frequency, by approximately 0.5 Hz from preschool age to age 6 or 7, confirming that mu frequency increases gradually throughout childhood, possibly reflecting changes in myelination, synaptic pruning or other neurophysiological changes that continue throughout childhood and adolescence (Sowell et al. 2004, Shaw et al. 2008).

The most novel observation in our study was a very non-linear, step-like transition of movement-evoked gamma band activity observed at movement onset. Unlike the preschool age children, all children in the 6 to 8-year-old cohort demonstrated an adult-like movement gamma synchronization (gamma ERS) at movement onset, in the typical frequency range of 70 to 90 Hz reported in MEG studies of adults (Cheyne et al. 2008, Dalal et al. 2008, Muthukumaraswamy 2010). Even more striking are the developmental changes observed in the longitudinal data, where children who had a low frequency gamma response at an earlier age now show an adult high frequency gamma ERS, as shown in Figure 6. Moreover, children who did have a high motor gamma ERS at preschool age in Session 1, showed a gamma ERS response that was virtually unchanged in both frequency and amplitude, and these patterns replicated for both left and right hand movements. This suggests that high-frequency movement-evoked gamma ERS emerges for both dominant and non-dominant hand movements, anywhere between 3 and 6 years of age and remains stable in frequency and amplitude afterwards. This is consistent with studies in adults showing that motor gamma ERS duration and peak frequency varies across individuals, but is highly similar within individuals in both frequency and duration for left and right hand movements, and this remains stable over time (Cheyne and Ferrari 2013). The role of movement-evoked gamma ERS is unknown, although it can be observed at various levels of the motor system, including subcortical structures (Lalo et al. 2008, Jenkinson et al. 2013), and appears to be related to higher cognitive aspects of movement execution such as movement awareness or effort, rather than specific movement parameters or sensory feedback (Muthukumaraswamy 2010, Cheyne and Ferrari 2013, Jenkinson et al. 2013).

### 4.3 Theoretical considerations and future studies

The physiological mechanisms underlying the developmental changes in brain activity observed in the current study are unknown. One currently popular hypothesis is that age-related changes in cortical oscillations are related to increases in the available pool of the inhibitory neurotransmitter GABA (Muthukumaraswamy et al. 2009, Gaetz et al. 2011). However, this is based on indirect evidence from a small number of MRI spectroscopy studies with inconsistent findings (Robson et al. 2015). Others have attributed age-related changes in cortical oscillations to altered long-range connectivity (Brookes et al. 2018). Although changes in local synaptic inputs and/or inter-areal connectivity likely play important roles in development, such hypotheses assume that the output organization of the primary motor cortex is relatively fixed from early childhood. The primary motor cortex and its associated efferent pathways has evolved across mammalian species in a way that is reflective of the need for increasing dexterity and fine motor control and is most advanced in non-human primates and humans (Lemon 2008). A major factor in this evolutionary trend is an increased role of the corticomotoneuronal (CM) system – descending pathways from primary motor cortex (MI) with monosynaptic connections directly onto spinal motor neurons – in the control of fine motor skills, particularly dexterous hand movements (Armand et al. 1997). This system has been shown to develop postnatally over an extended period in non-human primates, indicating that this is a late developing system both phylogenetically and ontogenetically, although its development in humans is unknown. Interestingly, compared to polysynaptic MI projections from the precentral gyrus to the spinal cord that are present at birth, CM neurons are located in more caudal portions of MI within the anterior wall of the central sulcus (Rathelot and Strick 2009). Thus, an alternative explanation for the dramatic increase in amplitude of MEG activity in motor cortex with age might be an increase in activity of neural populations of sulcal neurons in MI, as the CM system comes “on-line” along with increased motor skills during development.

A recent MEG study in children and adolescents from 9 to 15 years of age (Trevarrow et al. 2018) reported maturational changes in movement-related beta and gamma oscillations that peaked in strength in adolescence, followed by decreases in early adulthood. Relevant to the current study, their youngest participants showed movement-related cortical oscillatory activity similar in amplitude and timing to adults, suggesting that these motor networks may be relatively mature by 9 or 10 years of age. In contrast, our data show that movement-related brain activity is still undergoing changes after age 5 or 6 and it remains unclear at what age at which the transition to adult-like motor activity is complete, in particular, the emergence of the pre-movement readiness potential. Curiously, the absence of the readiness potential up to age 9 or 10 when general motor skills are relatively mature would suggest that this pre-movement brain activity does not play a direct role in motor execution. This is consistent however, with the general notion that role of MI in motor control is one of sensorimotor integration, such as the comparison of internal models and external feedback (corollary discharge), rather than sending specific motor commands to the muscles (Kalaska 2009) although such models have not been considered in the context of brain development.

Most importantly, our data confirm that there is a large degree of individual variability in the maturational trajectory of movement-related brain activity during early childhood. Since motor abilities were not assessed in the current study, it is unclear how the observed individual differences in movement-related brain activity may have been related to differences in motor abilities or other factors such as the development of hand dominance. Such studies will be particularly informative given that some of the youngest children in the study showed completely adult-like gamma ERS at less than 4 years of age, and more importantly, this remained stable at their second session, while other children develop this brain response much later. If MEG recordings of movement-related brain activity can be shown to track the rate of motor development across children, they may also provide useful indicators of atypical motor development, and whether motor disorders will resolve with age. Finally, our findings emphasize that individual variability in both the timing and frequency of movement-related oscillations, or other oscillatory responses needs be taken into account when studying differences in across fixed age groups.

## Acknowledgements

This work was supported by Australian Research Council grants DP170102407 and CE110001021. The authors acknowledge the collaboration of the Kanazawa Institute of Technology in establishing the KIT-Macquarie Brain Research Laboratory. C.J. and R.A. were supported by grants from the Canadian Natural Sciences and Engineering Research Council (DG184018-09) and the Canadian Institutes of Health Research.

